# Replacing H_2_O with D_2_O in cryo-EM reduces radiation damage in biological samples

**DOI:** 10.64898/2026.10.24.761413

**Authors:** Min Chevalier Kwon, Senik Matinyan, Eric van Genderen, Jan Pieter Abrahams

**Affiliations:** Biozentrum, University of Basel, 4056 Basel, Switzerland; Laboratory of Nanoscale Biology, Paul Scherrer Institut, 5232 Villigen PSI, Switzerland

## Abstract

Radiation damage limits the resolution achievable in electron microscopy of biological specimens. Hydrogen is the most abundant atom in these specimens, and its low mass maximizes the recoil energy it receives from incident electrons, making it the atom most readily lost under the beam. Substitution of hydrogen by deuterium halves this recoil energy and makes the bonds involved slightly harder to break, yet its protective effect on vitrified protein crystals has not been tested. Using cryogenic electron diffraction at 200 kV, we show that hen egg-white lysozyme crystals grown directly in D_2_O withstand a dose 1.7 times higher than crystals grown in H_2_O before their Bragg intensity at 10–7 Å falls to 1/*e* (13.2 ± 1.0 against 7.7 ± 0.4 e^−^/Å^2^), with every D_2_O-grown crystal outlasting every H_2_O-grown crystal. The protection is achieved with deuteration confined mainly to the solvent and the exchangeable hydrogens at the protein surface, which form the hydrogen bonds that hold the crystal lattice together. The protein itself need not be deuterated by expression in deuterated media. Direct crystallization in D_2_O involves only a single substitution in the mother liquor and offers an accessible strategy to extend the usable dose in electron diffraction of protein crystals.

## 1 Introduction

Electrons are the most effective probe of biological structure. For a given amount of radiation damage they yield more structural information than X-rays or neutrons, which is why cryo-electron microscopy (cryo-EM) and microcrystal electron diffraction (MicroED) can determine structures from isolated macromolecules and from crystals only a few hundred nanometres thick [1, 2]. The same interaction that makes electrons so informative also destroys the specimen. For every electron scattered elastically by a biological specimen, three to four are scattered inelastically, depositing a total of 60 to 80 eV that breaks bonds and creates radicals [1, 3]. In MicroED of proteinase K crystals, the attainable resolution dropped from 1.7 Å to 2.8 Å as the accumulated dose increased from 0.86 to 6 e^−^/Å^2^ [4]. Radiation damage therefore sets the dose that can be spent on each specimen, and with it the resolution that can be achieved and the smallest specimen that can be studied.

Much effort has gone into extending this limit. Cooling the specimen to liquid-nitrogen temperature has long been standard, as it suppresses the secondary chemistry that follows each ionization [3], and cooling further with liquid helium slows the fading of diffraction from two-dimensional crystals and has recently improved single-particle reconstructions [5, 6]. The energy of the incident electrons can also be chosen to maximize the information obtained per unit damage [7]. These approaches change the conditions of the experiment, the temperature or the beam, and leave the specimen itself unchanged.

The specimen offers another route. Hydrogen is the most abundant atom in proteins and in the water around them, and hydrogen gas is the main product of radiation damage in X-ray-irradiated protein crystals at cryogenic temperatures [8]. Under electron irradiation, the release of hydrogen is likewise the most common form of mass loss in frozen hydrated specimens [9]. Replacing hydrogen with deuterium doubles the nuclear mass while hardly changing the chemistry, and because the energy transferred from a beam electron to a nucleus depends on the mass of the nucleus [3, 9], isotope substitution could reduce damage without any change to the microscope. In transmission electron microscopy (TEM) at room temperature, deuteration has extended the lifetime of organic molecules encapsulated in carbon nanotubes [10] and of macromolecules imaged in D_2_O in graphene liquid cells (GLCs) [11].

For protein crystals, however, the evidence is negative or missing. In X-ray crystallography of lysozyme at 100 K, D_2_O gave no detectable protection [12], and lysozyme crystals examined by electron diffraction in GLCs at room temperature showed no significant difference between H_2_O and D_2_O [13]. To our knowledge, whether deuteration protects vitrified protein crystals under the cryogenic conditions used for structure determination by electron diffraction has not been measured. Such a measurement also depends on how the deuterium is introduced. Deuterating lysozyme crystals after they have grown leaves about twice as many amide hydrogens unexchanged as exchanging the protein before crystallization [14].

Here we grew lysozyme microcrystals directly in D_2_O-based mother liquor, so that the solvent was deuterated throughout crystal growth and the exchangeable hydrogens of the protein could exchange before the crystals formed, and compared them with crystals grown in H_2_O. We followed the decay of their Bragg intensity by cryogenic electron diffraction under continuous exposure at 200 kV, holding each crystal stationary and recording every frame with a hybrid pixel detector. Crystals grown in D_2_O withstood a dose 1.7 times higher before their Bragg intensity at 10–7 Å fell to 1/*e*.

## 2 Materials and methods

### 2.1 Lysozyme crystallization

Hen egg-white lysozyme (Sigma-Aldrich, 62970) was crystallized by the batch method at room temperature using protein concentrations of 25 mg/mL and 30 mg/mL in 0.7 M NaCl and 20 mM sodium acetate (pH 4.5). H_2_O-grown and D_2_O-grown crystals were prepared under otherwise identical conditions, with the solvent used for solution preparation as the sole intended variable. For H_2_O-grown crystals, lyophilized lysozyme was dissolved in crystallization solution prepared with ultrapure water (Milli-Q). For D_2_O-grown crystals, NaCl and sodium acetate were dissolved at the same nominal concentrations in deuterium oxide (Sigma-Aldrich, 151882, 99.9 atom% D), followed by dissolution of the lyophilized protein. This procedure maintained a deuterated solvent environment throughout crystal growth and permitted H/D exchange at exchangeable protein sites. Crystals formed under both conditions within two to three days.

### 2.2 Sample preparation for electron diffraction

Lacey carbon grids (200 mesh, Electron Microscopy Sciences) were glow-discharged for 45 s in air plasma (Quorum GloQube Plus). The crystal suspension was fragmented by repeated aspiration through a pipette tip. Grids were vitrified with a Leica EM GP plunge freezer at 20 ^°^C and 80 % relative humidity. During this procedure, 3 µL of the fragmented suspension was applied to the carbon side of the grid, and 0.5 µL of the corresponding buffer to the reverse side. Excess liquid was removed by sensor-controlled blotting for 5 s, and the grid was plunged into liquid ethane cooled by liquid nitrogen. Vitrified crystals were inspected on a Talos L120C microscope at 17,500× magnification and were about 1 µm across in both conditions (Figure 1a).

**Figure 1.**
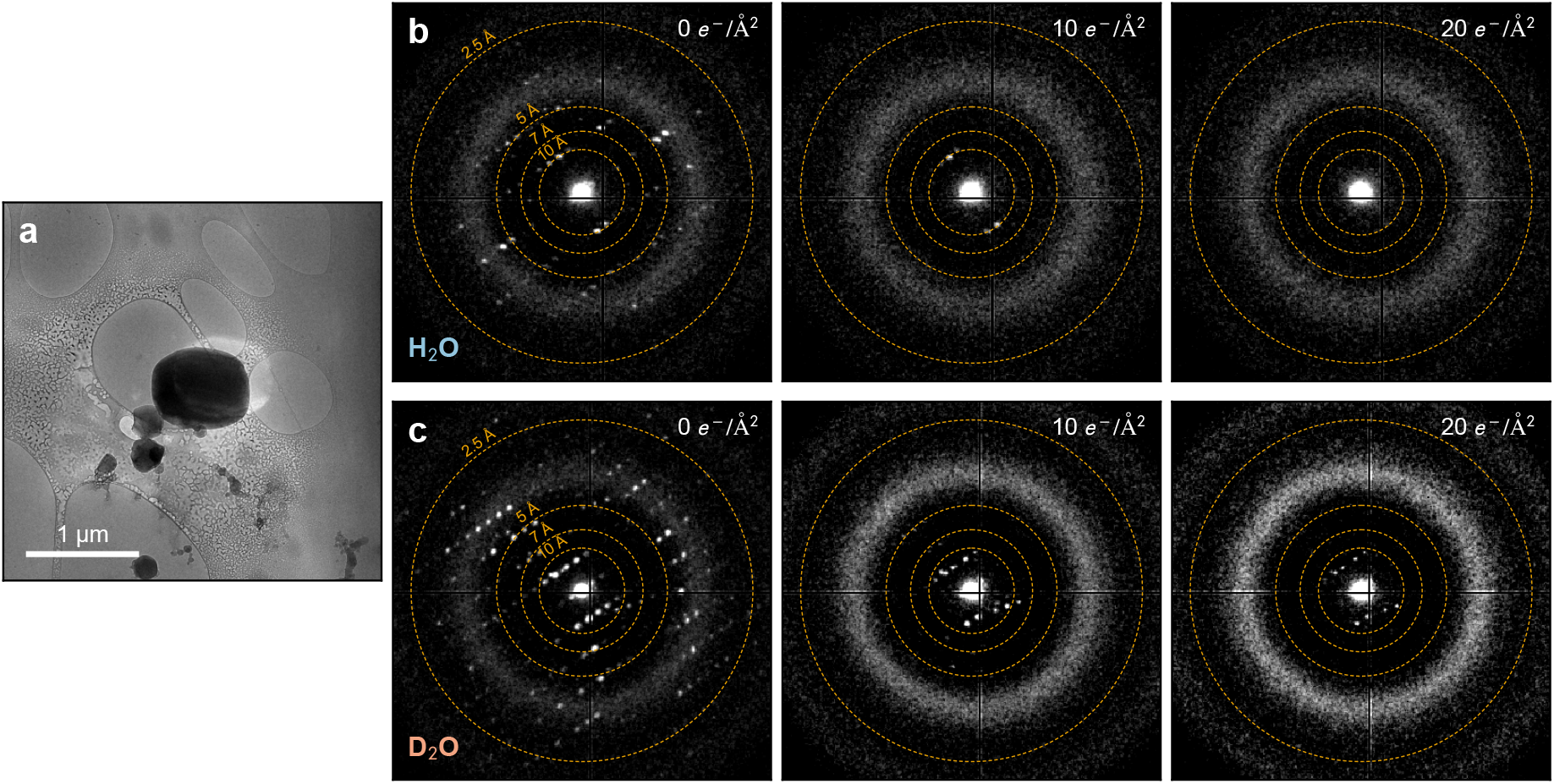
Lysozyme microcrystals and the decay of their electron diffraction. (a) Lysozyme microcrystal in vitreous ice, imaged during grid inspection. The beam used for diffraction was about 1.5 µm in diameter. (b, c) Diffraction patterns of an H_2_O-grown (b) and a D_2_O-grown (c) crystal at 0 e^−^/Å^2^, 10 e^−^/Å^2^ and 20 e^−^/Å^2^ . Each pattern is a single 30 ms frame (0.053 e^−^/Å^2^) shown on a square-root intensity scale. Dashed circles mark 10 Å, 7 Å, 5 Å and 2.5 Å, the boundaries of the resolution ranges in Figure 3.

### 2.3 Electron diffraction data collection

Diffraction patterns were recorded on a JEOL JEM-F200 TEM with a Schottky field emission gun operated at 200 kV, using an ASI Cheetah M3 hybrid pixel detector (Medipix3, 55 µm pixels). Specimens were kept at liquid-nitrogen temperature in a Gatan ELSA cryo-transfer holder. Parallel illumination was set with spot size 5, alpha 3 and a 30 µm condenser aperture, giving a beam around 1.5 µm in diameter at the specimen. And the camera length, calibrated with a polycrystalline aluminium standard, was 820 mm. The dose rate, measured with the detector in vacuum over the calibrated beam area, was 1.76 e^−^/Å^2^/s, corresponding to 0.053 e^−^/Å^2^ per 30 ms frame. The base brightness of the gun was reduced to reach this dose rate.

Crystals were located in diffraction mode while the detector recorded continuously. Once Bragg reflections appeared, the stage was stopped and the stationary crystal was exposed continuously for up to about 50 e^−^/Å^2^ (Figure 1b,c). Dose was counted from the first frame in which the crystal was fully illuminated. Four H_2_O-grown and four D_2_O-grown crystals were measured.

### 2.4 Extraction of Bragg intensities

Bragg intensities were separated from the diffuse background of each frame with *radialBragg*, a program developed in our laboratory (Figure 2). The program locates the direct beam by optimizing the two-fold rotational symmetry of the pattern at low spatial frequencies and transforms the frame to polar coordinates around it. In each radial bin, the background is estimated as the median pixel value after iterative rejection of outliers more than three standard deviations above the mean, and its variance from the remaining pixels. Each pixel is then expressed as a signal-to-noise ratio (SNR), the number of standard deviations by which it exceeds the background of its radial bin (Figure 2b). Connected pixels above a seed threshold mark the strongest reflections, and an auto-convolution at their positions gives the lattice periodicity. Further pixels are accepted if they lie at lattice-commensurate positions and exceed a lower threshold, and isolated single pixels are rejected. The background-subtracted values of the masked pixels (Figure 2c) are summed in each radial bin with fractional pixel weights, giving the integrated Bragg intensity as a function of resolution (Figure 2d).

**Figure 2.**
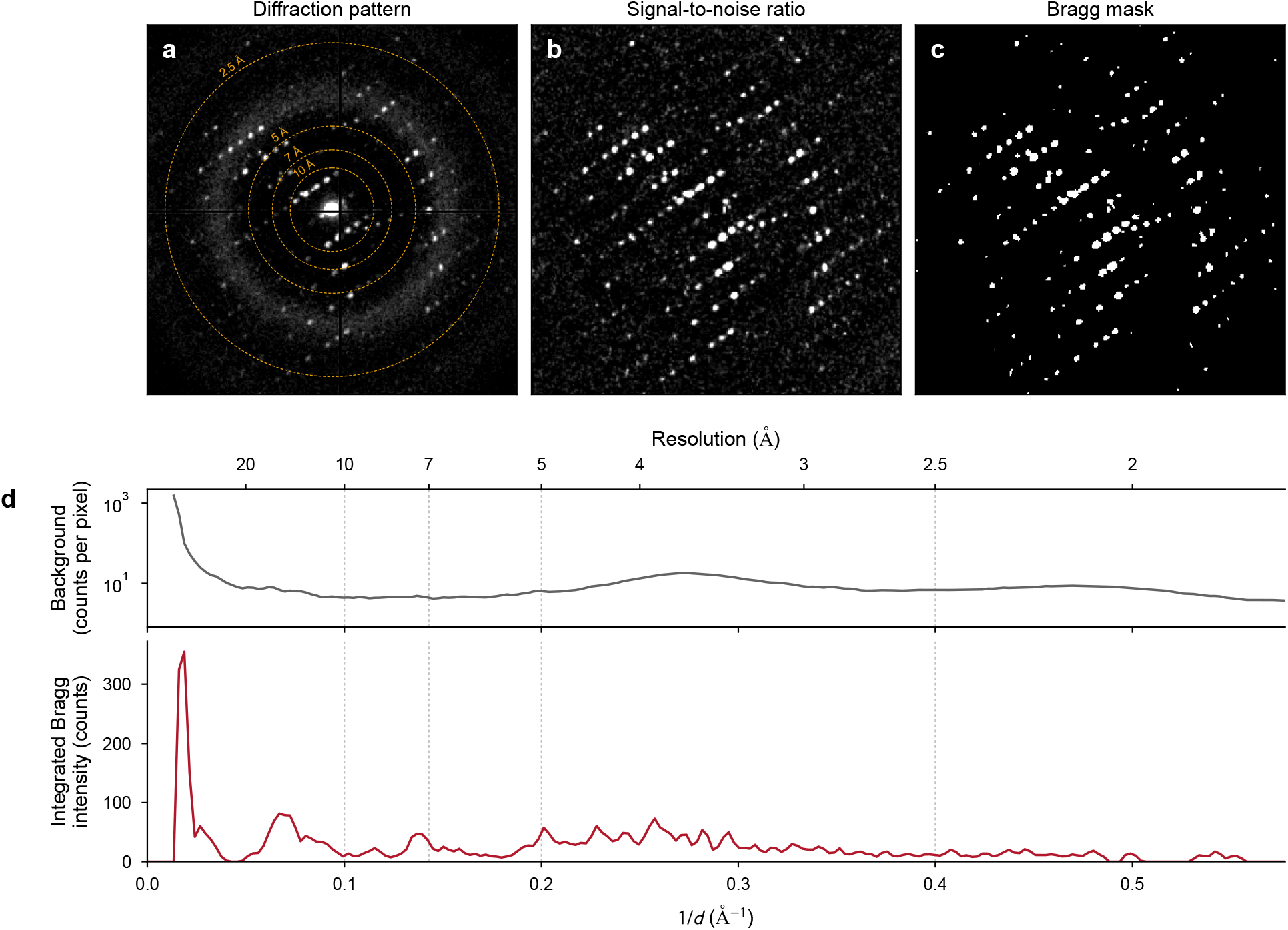
Extraction of Bragg intensities with *radialBragg*, shown for the first frame of a D_2_O-grown crystal. A weaker second lattice from another D_2_O-grown crystal in the beam is also visible. (a) Diffraction pattern with resolution circles. (b) SNR of each pixel relative to the background of its radial bin. (c) Bragg mask. (d) Radial profiles of the background (top, counts per pixel, logarithmic scale) and of the background-subtracted Bragg intensity integrated over each one-pixel ring (bottom). The broad maximum of the background near 3.7 Å arises from vitreous ice. Dotted lines mark the resolution ranges of Figure 3.

### 2.5 Characteristic dose

For each frame, the integrated Bragg intensity was summed over three resolution ranges, 10–7 Å, 7–5 Å and 5–2.5 Å, averaged over blocks of ten consecutive frames (0.53 e^−^/Å^2^ per ten frames) and normalized to the first block. The characteristic dose *D*_*e*_ was defined as the dose at which the normalized intensity of a crystal fell to 1/*e*, obtained by linear interpolation between blocks. *D*_*e*_ is reported as mean ± standard error of the mean of the four crystals per condition, and the isotope effect as the ratio of the mean *D*_*e*_ of D_2_O-grown to H_2_O-grown crystals. Data were analysed in Python.

## 3 Results

Crystals of both conditions diffracted to 2.5 Å or better in the first frames (Figure 1b,c). In the radial profiles, the Bragg intensity lay on a diffuse background with a broad maximum near 3.7 Å from vitreous ice (Figure 2d). Under continuous exposure, the reflections faded first at high resolution. At 10 e^−^/Å^2^, reflections beyond about 5 Å had disappeared in both conditions, whereas low-resolution reflections remained more prominent in the D_2_O-grown crystal (Figure 1b,c).

In both conditions, the Bragg intensity decayed faster at higher resolution (Figure 3, Table 1). In the 10–7 Å range, the intensity of H_2_O-grown crystals declined from about 2 e^−^/Å^2^, whereas that of D_2_O-grown crystals remained near its initial value for about 5 e^−^/Å^2^ before declining. D_2_O-grown crystals reached 1/*e* at a 1.72 times higher dose, with *D*_*e*_ of 13.2 ± 1.0 against 7.7 ± 0.4 e^−^/Å^2^, and each of the four D_2_O-grown crystals (10.9–15.9 e^−^/Å^2^) exceeded each of the four H_2_O-grown crystals (6.5–8.6 e^−^/Å^2^). In the 7–5 Å range, the curves of the two conditions coincided up to about 5 e^−^/Å^2^, after which D_2_O-grown crystals decayed more slowly (ratio 1.23), although the values of individual crystals overlapped. In the 5–2.5 Å range, the intensity declined from the first frames and the curves of the two conditions nearly coincided (ratio 0.85).

**Table 1.** Characteristic dose *D*_*e*_ of H_2_O-grown (H_2_O) and D_2_O-grown (D_2_O) crystals. Values are mean ± standard error of four crystals per condition, with the range of individual crystals in parentheses. Ratio, *D*_*e*_(D_2_O)/*D*_*e*_(H_2_O).

| Resolution ($\text{\AA}$) | $D_e \text{ (e}^-/\text{\AA}^2\text{)}$ | | Ratio |
| --- | --- | --- | --- |
| | $\text{H}_2\text{O}$ | $\text{D}_2\text{O}$ | |
| 10–7 | $7.7 \pm 0.4$ (6.5–8.6) | $13.2 \pm 1.0$ (10.9–15.9) | 1.72 |
| 7–5 | $6.8 \pm 1.1$ (3.8–8.9) | $8.3 \pm 1.2$ (5.9–10.5) | 1.23 |
| 5–2.5 | $5.1 \pm 0.5$ (4.5–6.7) | $4.3 \pm 0.6$ (3.1–5.5) | 0.85 |

**Figure 3.**
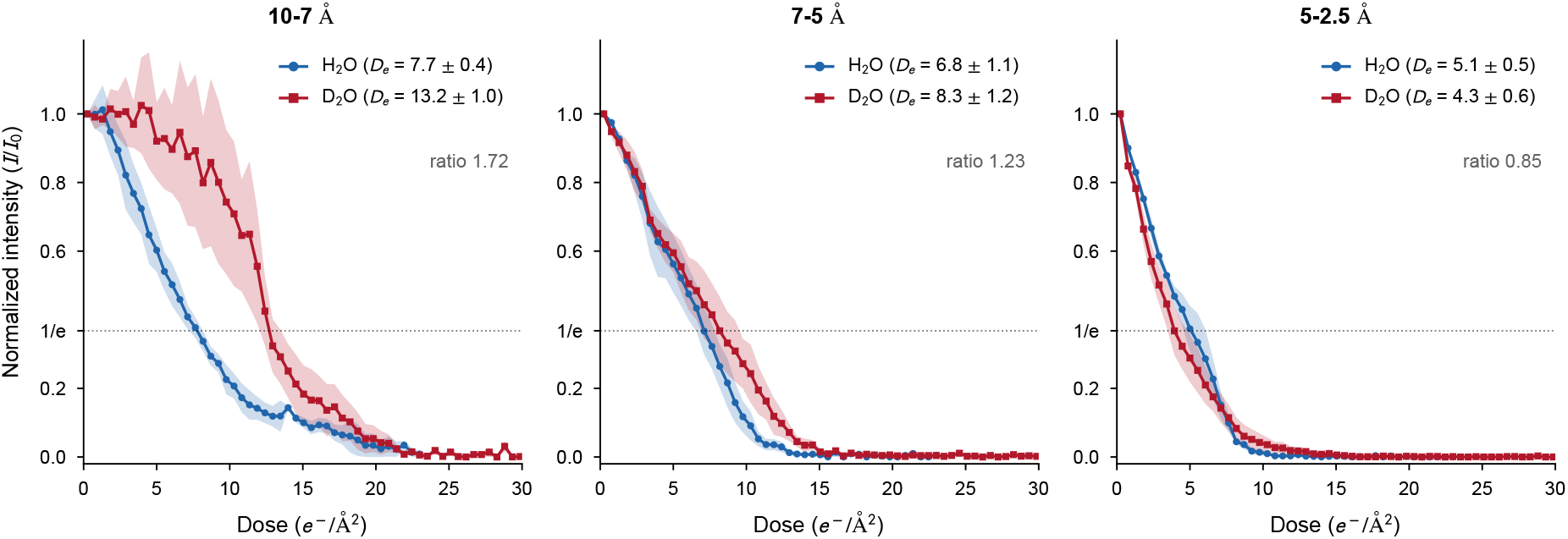
Decay of the Bragg intensity with dose for H_2_O-grown (H_2_O, blue) and D_2_O-grown (D_2_O, red) crystals in three resolution ranges. Intensities were averaged over blocks of ten frames (0.53 e^−^/Å^2^ per ten frames) and normalized to the first block. Lines and markers show the mean of four crystals per condition and shaded bands the standard error of the mean. *D*_*e*_ is given as mean ± standard error of the four crystals, and the ratio as *D*_*e*_(D_2_O)/*D*_*e*_(H_2_O). In the 10–7/Å range, the wide D_2_O band over the first few e^−^ Å^2^ arises from one crystal with few reflections in this range, whose reflections entered and left the diffraction condition in individual frames.

In stationary diffraction, the diffracted intensity depends on crystal size and orientation. To test whether diffraction strength could account for the difference between the conditions, we compared *D*_*e*_ with the initial Bragg intensity of each crystal (Figure 4). Within each condition, crystals with stronger initial diffraction tended to have larger *D*_*e*_. The initial intensities of the two conditions spanned the same range, from 1.2 to 2.8 × 10^3^ counts for H_2_O-grown and from 0.9 to 3.1 × 10^3^ counts for D_2_O-grown crystals, and the most weakly diffracting D_2_O-grown crystal had a larger *D*_*e*_ than the most strongly diffracting H_2_O-grown crystal (12.9 against 8.6 e^−^/Å^2^). The difference between the conditions is therefore not explained by diffraction strength.

**Figure 4.**
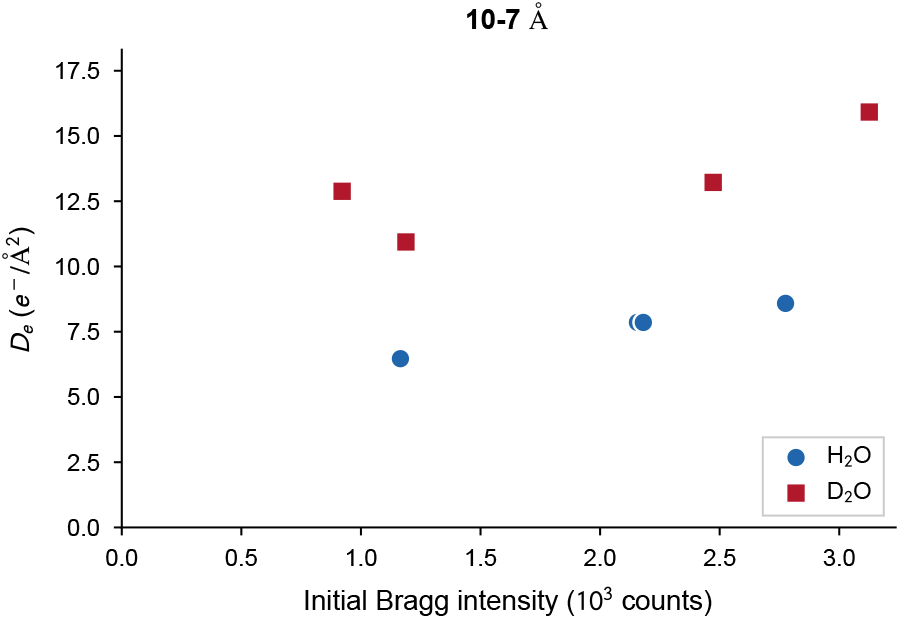
Characteristic dose in the 10–7 Å range of each crystal against its initial Bragg intensity, summed over 10–2.5 Å in the first ten frames. H_2_O and D_2_O denote H_2_O-grown and D_2_O-grown crystals. Two H_2_O points overlap.

## 4 Discussion

### 4.1 Isotope effect under cryogenic conditions

Growing lysozyme crystals in D_2_O increased their tolerance to the electron dose by a factor of 1.7 at 10–7 Å (Table 1). Isotope effects of comparable magnitude have been reported in electron microscopy at room temperature. An isotope effect of comparable magnitude has been reported in electron microscopy at room temperature, where deuteration extended the lifetime of organic molecules encapsulated in carbon nanotubes by about a factor of two [10]. The earlier measurements on lysozyme crystals that found no protection also grew the crystals in D_2_O but used X-rays [12] or crystals in liquid at room temperature [13]. Our electron diffraction measurements show that the isotope effect is present in protein crystals under cryogenic conditions.

### 4.2 Origin of the protection

One reason why D_2_O may be more resistant to electron irradiation is that deuterium is heavier than hydrogen. In an elastic collision, the incident electron transfers momentum *q* to a nucleus. The resulting recoil energy is

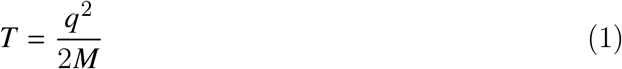

where *M* is the nuclear mass. A deuterium nucleus is about twice as heavy as a hydrogen nucleus, so for the same momentum transfer it receives only about half the recoil energy, 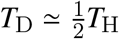. Its recoil velocity is also smaller, *v*_D_ ≃ *v*_H_/2. The heavier D nucleus is simply harder to move. At 200 kV, the maximum recoil energy is about 520 eV for hydrogen and 260 eV for deuterium. If the recoil energy is sufficiently large, it will cause bond breakage [9]. At lower energy, the displaced nuclei are pulled back towards equilibrium, and the recoil energy appears as bond vibration.

The effect of the larger deuterium mass may therefore not be limited to collisions energetic enough to cause immediate knock-on displacement. The size of the nuclear excursion following a sub-threshold collision can be estimated by treating the bond as a harmonic oscillator with force constant *k*. As the recoiling nucleus moves away from its equilibrium position, its recoil energy is converted into potential energy of the bond. At the maximum displacement *A*, this gives

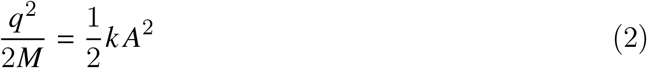

where *A* is the resulting nuclear excursion from equilibrium. Thus,

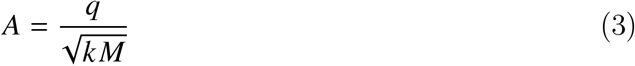

For the same momentum transfer, replacement of H by D therefore reduces the amplitude of the induced nuclear motion by approximately 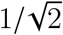. The heavier D nucleus is displaced less far from its equilibrium position, producing a smaller distortion of the bond and making it less likely to stretch the bond far enough for dissociation to occur.

There is another effect that works in the same direction. Although replacing H by D hardly changes the electronic potential of an O–H bond, it does change its vibrational energy levels. In the harmonic approximation,

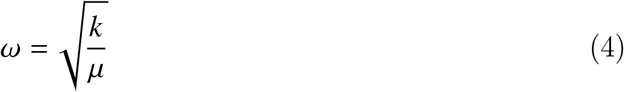

where *k* is the bond force constant and *µ* its reduced mass. The larger reduced mass of O–D gives it a lower vibrational frequency and, with 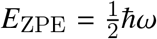, a lower zero-point energy. This isotope shift is well established spectroscopically. For gas-phase HDO, the O–H and O–D stretching frequencies are approximately 3707 cm^−1^ and 2727 cm^−1^, respectively [15]. The bottom of the O–D vibrational ground state therefore lies a little further below the dissociation limit than that of O–H. In this sense the O–D bond is slightly harder to break, even though the underlying electronic bond is essentially unchanged. Experimentally, dissociation energies of about 5.19 eV for HO–H and 5.28 eV for DO–D have been reported [16].

Thus, when an electron gives the system a kick, deuteration helps in two ways. The heavier nucleus takes up less recoil energy for a given momentum transfer, and slightly more energy is required to break the bond. In D_2_O-grown crystals, both effects reduce the disturbance of the bonds to deuterium for a given dose, consistent with the slower decay of the Bragg intensity that we observe.

### 4.3 Distribution of deuterium in the crystal

At high resolution, H_2_O-grown and D_2_O-grown crystals decayed alike (Table 1), which can be traced to where the deuterium resides in the crystal. Crystallization in D_2_O replaces the solvent and the exchangeable hydrogens of the protein, those bound to nitrogen and oxygen, whereas hydrogens bound to carbon remain protiated. In a native protein exposed to D_2_O, hydrogens in the core also exchange incompletely, so that only 10–20% of all its hydrogen atoms are replaced by deuterium [14]. Neutron crystallography of tetragonal lysozyme that was exchanged in D_2_O before crystallization, as in this work, shows that even after several months the amide hydrogens forming main-chain hydrogen bonds in helices and *β*-sheets remain unexchanged, whereas most other amide hydrogens are fully or partially exchanged [14]. Exchange at these positions requires unfolding and refolding of the protein [17]. Deuterium in D_2_O-grown crystals is therefore concentrated in the solvent and in the hydrogen bonds at the protein surface, while the hydrogen bonds that hold the helices and *β*-sheets together keep their hydrogens.

The mass effects described above act only on bonds to deuterium and matter most for weak interactions. Recoil energies and nuclear excursions that leave a covalent O–H bond intact can still break or distort a hydrogen bond, which is more than an order of magnitude weaker. In D_2_O-grown crystals, deuterium sits in the hydrogen bonds of the solvent and of the protein surface, including the contacts between molecules that hold the lattice together.

In the Debye–Waller model applied to radiation damage, the intensity of a reflection at resolution *d* scales as *I*/*I*_0_ = exp(−*B*/2*d*^2^), where *B* = 8*π*^2^⟨*u*^2^⟩ and ⟨*u*^2^⟩ is the mean-square displacement of the atoms. It follows that the intensity drops to 1/*e* when the root-mean-square displacement reaches *d*/2*π*. This threshold is about 0.4 Å at 2.5 Å resolution but increases to 1.6 Å at 10 Å. Small displacements from damage throughout the largely protiated molecule therefore remove the high-resolution reflections, whereas reflections at 10–7 Å persist until molecules shift or rotate as the lattice contacts break. Deuteration of these contacts slows the loss of lattice order and thus the decay at 10–7 Å, while the decay at high resolution changes little.

Perdeuterating proteins by expressing them in deuterated media is common practice in NMR studies [18]. Assuming that the protective mechanism we proposed also applies to H/D atoms located in the molecular core, we anticipate such perdeuterated samples will have an increased radiation tolerance in single-particle cryo-EM, improving the quality of such studies.

## 5 Conclusions

We have shown that lysozyme crystals grown in D_2_O withstand a dose 1.7 times higher than crystals grown in H_2_O before their Bragg intensity at 10–7 Å falls to 1/*e* in cryogenic electron diffraction. The protection is consistent with the larger mass of deuterium, which takes up less recoil energy in elastic collisions and forms bonds that are slightly harder to break, and it acts mainly on the deuterated hydrogen bonds of the solvent and the protein surface that hold the crystal lattice together.

Unlike perdeuteration, which requires the protein to be expressed in deuterated media, growing crystals in D_2_O only replaces the water used for crystallization, needs a few millilitres of D_2_O and leaves the crystallization and vitrification protocols unchanged. It therefore offers a simple way to slow radiation damage in electron diffraction of protein crystals. Because the isotope effect acts on the nuclei, it is likely to be complementary to other strategies for reducing radiation damage, such as the choice of accelerating voltage [7] or cooling to liquid-helium temperature [5, 6]. Combining these approaches, and extending the measurements to more crystals and other proteins, may further increase the dose that biological specimens tolerate in cryo-EM and diffraction.

### Declaration of generative AI and AI-assisted technologies in the manuscript preparation process

During the preparation of this work the authors used Claude (Anthropic) and ChatGPT (OpenAI) in order to improve the readability of the manuscript and to assist with coding. After using these tools, the authors reviewed and edited the content as needed and take full responsibility for the content of the published article.

## References

[1] R. Henderson, The potential and limitations of neutrons, electrons and X-rays for atomic resolution microscopy of unstained biological molecules, Quarterly Reviews of Biophysics 28 (2) (1995) 171–193. doi:10.1017/S003358350000305X.

[2] D. Shi, B. L. Nannenga, M. G. Iadanza, T. Gonen, Three-dimensional electron crystallography of protein microcrystals, eLife 2 (2013) e01345. doi:10.7554/eLife.01345.

[3] M. C. Kwon, J. P. Abrahams, Elastic and inelastic interactions of electrons in transmission electron microscopy, Ultramicroscopy 284 (2026) 114383. doi:10.1016/j.ultramic.2026.114383.

[4] J. Hattne, D. Shi, C. Glynn, C.-T. Zee, M. Gallagher-Jones, M. W. Martynowycz, J. A. Rodriguez, T. Gonen, Analysis of Global and Site-Specific Radiation Damage in Cryo-EM, Structure 26 (5) (2018) 759–766.e4. doi:10.1016/j.str.2018.03.021.

[5] K. Naydenova, A. Kamegawa, M. J. Peet, R. Henderson, Y. Fujiyoshi, C. J. Russo, On the reduction in the effects of radiation damage to two-dimensional crystals of organic and biological molecules at liquid-helium temperature, Ultramicroscopy 237 (2022) 113512. doi:10.1016/j.ultramic.2022.113512.

[6] J. L. Dickerson, K. Naydenova, M. J. Peet, H. Wilson, B. Nandy, G. McMullan, R. Morrison, C. J. Russo, Reducing the effects of radiation damage in cryo-EM using liquid helium temperatures, Proceedings of the National Academy of Sciences 122 (17) (2025) e2421538122. doi:10.1073/pnas.2421538122.

[7] M. J. Peet, R. Henderson, C. J. Russo, The energy dependence of contrast and damage in electron cryomicroscopy of biological molecules, Ultramicroscopy 203 (2019) 125–131. doi:10.1016/j.ultramic.2019.02.007.

[8] A. Meents, S. Gutmann, A. Wagner, C. Schulze-Briese, Origin and temperature dependence of radiation damage in biological samples at cryogenic temperatures, Proceedings of the National Academy of Sciences 107 (3) (2010) 1094–1099. doi: 10.1073/pnas.0905481107.

[9] R. F. Egerton, Radiation damage to organic and inorganic specimens in the TEM, Micron 119 (2019) 72–87. doi:10.1016/j.micron.2019.01.005.

[10] T. W. Chamberlain, J. Biskupek, S. T. Skowron, P. A. Bayliss, E. Bichoutskaia, U. Kaiser, A. N. Khlobystov, Isotope Substitution Extends the Lifetime of Organic Molecules in Transmission Electron Microscopy, Small 11 (5) (2015) 622–629. doi: 10.1002/smll.201402081.

[11] H. Wang, K. H. Nagamanasa, Y.-J. Kim, O.-H. Kwon, S. Granick, Longer-Lasting Electron-Based Microscopy of Single Molecules in Aqueous Medium, ACS Nano 12 (8) (2018) 8572–8578. doi:10.1021/acsnano.8b04190.

[12] J. Kmetko, M. Warkentin, U. Englich, R. E. Thorne, Can radiation damage to protein crystals be reduced using small-molecule compounds?, Acta Crystallographica Section D Biological Crystallography 67 (10) (2011) 881–893. doi: 10.1107/S0907444911032835.

[13] S. Plana-Ruiz, A. Gómez-Pérez, M. Budayova-Spano, D. L. Foley, J. Portillo-Serra, E. Rauch, E. Grivas, D. Housset, P. P. Das, M. L. Taheri, S. Nicolopoulos, W. L. Ling, High-Resolution Electron Diffraction of Hydrated Protein Crystals at Room Temperature, ACS Nano (Oct. 2023). doi:10.1021/acsnano.3c05378.

[14] A. Kita, Y. Morimoto, Hydrogen/deuterium exchange behavior in tetragonal hen egg-white lysozyme crystals affected by solution state, Journal of Applied Crystallography 53 (Pt 3) (2020) 837–840. doi:10.1107/S1600576720005488.

[15] T. Shimanouchi, Tables of molecular vibrational frequencies, consolidated volume i:, Tech. Rep. NBS NSRDS 39, National Institute of Standards and Technology, Gaithersburg, MD (Jan. 1972). doi:10.6028/NBS.NSRDS.39.

[16] S. Kumar, M. Hoshino, B. Kerkeni, G. García, P. Limão-Vieira, Correction to “Isotope Effect in D2 O Negative Ion Formation in Electron Transfer Experiments: DO–D Bond Dissociation Energy”, The Journal of Physical Chemistry Letters 14 (40) (2023) 9002–9002. doi:10.1021/acs.jpclett.3c02406.

[17] A. Kita, Y. Morimoto, An Effective Deuterium Exchange Method for Neutron Crystal Structure Analysis with Unfolding-Refolding Processes, Molecular Biotechnology 58 (2) (2016) 130–136. doi:10.1007/s12033-015-9908-8.

[18] K. H. Gardner, L. E. Kay, The Use of 2h, 13c, 15n Multidimensional Nmr Gto Study the Structure and Dynamics of Proteins, Annual Review of Biophysics 27 (1998) 357–406. doi:10.1146/annurev.biophys.27.1.357.

