## Supplementary Figure and Table for "Replacing H_2_O with D_2_O in cryo-EM reduces radiation damage in biological samples"

### A Supplementary Figure and Table

*radialBragg* identifies Bragg pixels with a seed threshold and a lower threshold on the SNR, as described in the main text. The default values of 5 and 3 admit more weak pixels into the mask than the values of 6 and 4, and the masks obtained with 6 and 4 follow the reflections visible in the diffraction patterns more closely (Figure A1). In frames recorded after the reflections had visibly faded, the default thresholds still assigned background pixels to the mask, which added up to 12% of the initial Bragg intensity, whereas the stricter thresholds gave empty masks. Thresholds of 6 and 4 were therefore used throughout this work. Weaker signal falls below stricter thresholds at a lower dose, and the default thresholds gave higher values of $\boldsymbol{D_e}$ in all three resolution ranges (Table A1). The comparison between the two conditions was nevertheless the same with both settings. At 10–7 Å, $D_2O$-grown crystals withstood a dose 1.6 to 1.7 times higher, with every $D_2O$-grown crystal exceeding every $H_2O$-grown crystal, and at 5–2.5 Å the two conditions did not differ.

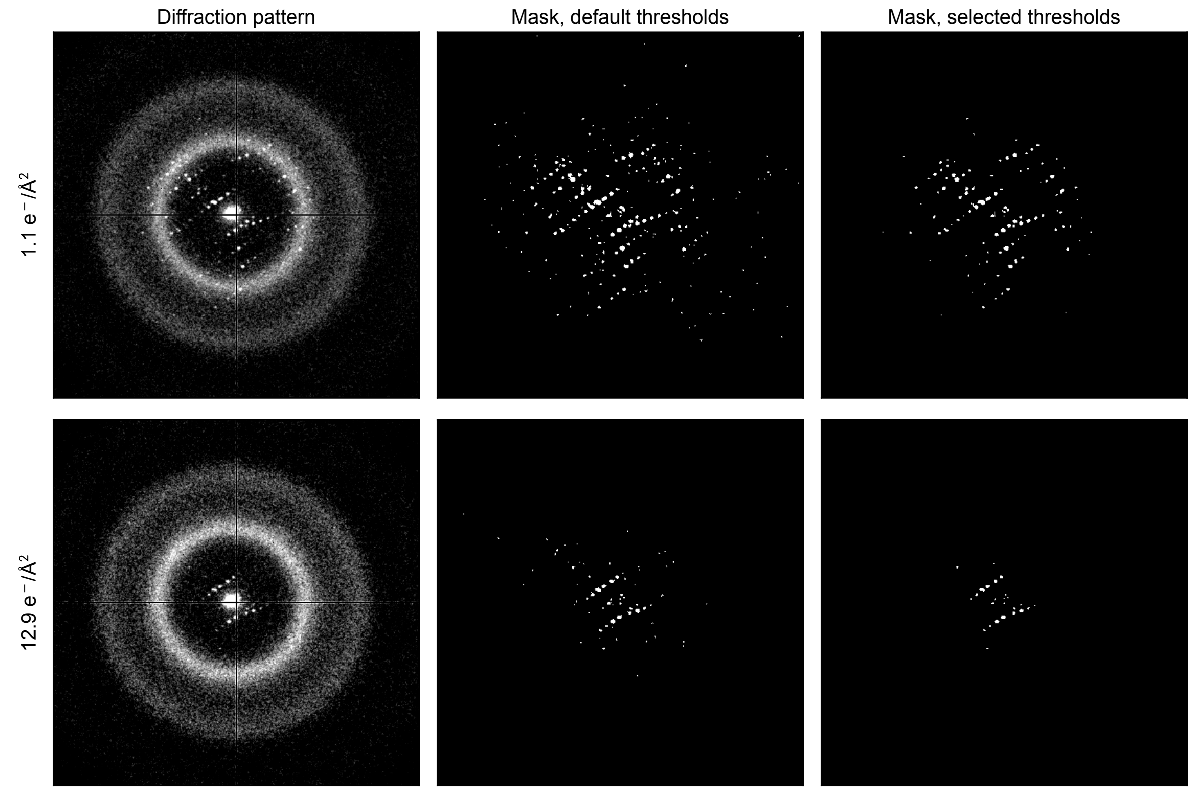


**Figure A1:** Bragg masks of a $D_2O$-grown crystal obtained with the default seed and lower thresholds of *radialBragg*, 5 and 3 (middle), and with the thresholds used in this work, 6 and 4 (right), together with the diffraction patterns (left), after $1.1\,e^-/\text{Å}^2$ (top) and $12.9\,e^-/\text{Å}^2$ (bottom). The default thresholds admit more weak pixels at both doses, whereas the masks obtained with the stricter thresholds follow the visible reflections more closely.

**Table A1:** Characteristic dose $\boldsymbol{D_e}$ ($e^-/\text{Å}^2$) obtained with the detection thresholds used in this work and with the default thresholds of *radialBragg*. Values are mean ± standard error of four crystals per condition. Ratio, $\boldsymbol{D_e}(D_2O)/\boldsymbol{D_e}(H_2O)$.

| | Seed 6, lower 4 | | | Seed 5, lower 3 | | |
|---|---|---|---|---|---|---|
| Resolution (Å) | $H_2O$ | $D_2O$ | Ratio | $H_2O$ | $D_2O$ | Ratio |
| 10–7 | 7.7 ± 0.4 | 13.2 ± 1.0 | 1.72 | 9.7 ± 0.9 | 15.2 ± 0.7 | 1.57 |
| 7–5 | 6.8 ± 1.1 | 8.3 ± 1.2 | 1.23 | 8.6 ± 0.5 | 11.2 ± 1.8 | 1.30 |
| 5–2.5 | 5.1 ± 0.5 | 4.3 ± 0.6 | 0.85 | 6.8 ± 0.4 | 6.9 ± 0.7 | 1.01 |
